# SARS-CoV-2 Spike protein activates TMEM16F-mediated platelet pro-coagulant activity

**DOI:** 10.1101/2021.12.14.472668

**Authors:** Ambra Cappelletto, Harriet E. Allan, Marilena Crescente, Edoardo Schneider, Rossana Bussani, Hashim Ali, Ilaria Secco, Simone Vodret, Roberto Simeone, Luca Mascaretti, Serena Zacchigna, Timothy D. Warner, Mauro Giacca

## Abstract

**Background:** Thrombosis of the lung micro-vasculature is a characteristic of COVID-19 disease, which is observed in large excess compared to other forms of acute respiratory distress syndrome and thus suggests a trigger for thrombosis endogenous to the lung. Our recent work has shown that the SARS-CoV-2 Spike protein activates the cellular TMEM16F chloride channel and scramblase. Through a screening on >3,000 FDA/EMA approved drugs, we identified Niclosamide and Clofazimine as the most effective molecules at inhibiting this activity. As TMEM16F plays an important role in the stimulation of the pro-coagulant activity of platelets, and considering that platelet abnormalities are common in COVID-19 patients, we investigated whether Spike directly affects platelet activation and pro-thrombotic function and tested the effect of Niclosamide and Clofazimine on these processes.

**Methods:** We produced SARS-CoV-2 Spike or VSV-G protein-pseudotyped virions, or generated cells expressing Spike on their plasma membrane, and tested their effects on platelet adhesion (fluorescence), aggregation (absorbance), exposure of phosphatidylserine (flow cytometry for annexin V binding), calcium flux (flow cytometry for fluo-4 AM), and clot formation and retraction. These experiments were also conducted in the presence of the TMEM16F activity inhibitors Niclosamide and Clofazimine.

**Results:** Here we show that exposure to SARS-CoV-2 Spike promotes platelet activation, adhesion and spreading, both when present on the envelope of virions or upon expression on the plasma membrane of cells. Spike was effective both as a sole agonist or by enhancing the effect of known platelet activators, such as collagen and collagen-related peptide. In particular, Spike exerted a noticeable effect on the procoagulant phenotype of platelets, by enhancing calcium flux, phosphatidylserine externalisation, and thrombin generation. Eventually, this resulted in a striking increase in thrombin-induced clot formation and retraction. Both Niclosamide and Clofazimine almost abolished this Spike-induced pro-coagulant response.

**Conclusions:** Together, these findings provide a pathogenic mechanism to explain thrombosis associated to COVID-19 lung disease, by which Spike present in SARS-CoV-2 virions or exposed on the surface of infected cells, leads to local platelet stimulation and subsequent activation of the coagulation cascade. As platelet TMEM16F is central in this process, these findings reinforce the rationale of repurposing drugs targeting this protein, such as Niclosamide, for COVID-19 therapy.

## INTRODUCTION

Thrombosis is a defining characteristic of COVID-19 lung pathology in patients with severe disease. Recent meta-analyses of published studies reveal that clinical indicators of thrombosis, including elevated D-dimers, fibrinogen and thrombosis-associated inflammatory biomarkers, are present in over 20% of patients with COVID-19 and in at least half of patients requiring intensive care ^1–5^. At the histological level, our own post-mortem analysis of 41 patients with COVID-19 revealed the presence of microthrombosis in 83% of patients requiring intensive care and 75% of all the other patients ^6^. This high prevalence resonates with that of several other pathology investigations over the last year ^7–11^. While a thrombotic response is common in other respiratory infections and other causes of acute respiratory distress syndrome (ARDS), the magnitude of this response appears a characteristic of COVID-19. A post-mortem study reported that severe vascular injury, including alveolar microthrombi, was 9 times more prevalent in the lungs of COVID-19 patients than in patients with influenza ^7^.

At least three observations suggest that thrombosis in COVID-19 is triggered by local events occurring in the infected lungs. First, thrombi and fibrin deposition in the lungs are asynchronous, by which relatively recent thrombi infiltrated by inflammatory cells are close to older thrombi, in advanced stage of fibrotic organization ^6^. This is consistent with a pro-thrombotic predisposition endogenous to the infected lungs, as opposed to multiple embolic events. Second, the presence of macro- or micro-vascular thrombosis is only sporadically detected in other organs ^6–11^. Again, this is suggestive that viral infection triggers local thrombotic events in the lungs. Third, patients with severe COVID-19 have high levels of D-dimer (a product of fibrin degradation) and fibrinogen, but do not show an increase in prothrombin time or a decrease in antithrombin levels and rarely develop disseminated intravascular coagulation (DIC)^12^. This argues against thrombosis as a consequence of a systemic consumptive coagulopathy.

What the causes are of the endogenous predisposition to lung thrombosis remains debated. This pulmonary intravascular coagulopathy ^13^ has variously been attributed to endothelial dysfunction, possibly triggered by direct SARS-CoV-2 infection of endothelial cells ^6, 14^ or to the hyperinflammatory state that accompanies SARS-CoV-2 infection, sustained by various pro-inflammatory cytokines (in particular, IL-6, TNFα, and IL-1β) ^15–19^. A proposed mechanism connecting inflammation and thrombosis is immunothrombosis, by which neutrophils and monocytes activate the coagulation cascade as a host immune defence against infection ^20–22^. In the infected lungs, neutrophil extracellular traps (NETs) released by neutrophils may contribute to vascular occlusion and formation of thrombi^23, 24^.

While these mechanisms most likely contribute to the local pro-thrombotic state, they still do not explain why lung thrombosis is particularly frequent in COVID-19, compared to other causes of ARDS, while the whole concept of hyperinflammation and cytokine storm has been recently questioned ^25^.

Several observations point to a specific involvement of platelets in the pathogenesis of thrombosis in COVID-19 patients. A common characteristic of these patients is thrombocytopenia ^5, 26–28^. COVID-19 lungs show increased number of pulmonary megakaryocytes, which could be indicative of increased local megakaryopoiesis in response to platelet consumption ^29^. SARS-CoV-2 infection is associated with platelet hyperreactivity, including increased P-selectin expression, increased activation and spreading on both fibrinogen and collagen, and elevated levels of circulating platelet-neutrophil, -monocyte, and -T-cell aggregates ^30–32^. The actual molecular mechanisms leading to this platelet hyperactivity and consumption during SARS-CoV-2 infection, coupled with local thrombosis, remain largely elusive.

While studying the molecular properties of the SARS-CoV-2 Spike protein, we discovered a novel mechanism that regulates cell-cell fusion induced by this protein. We started from the observation that the lungs of almost 90% of COVID-19 patients contain several syncytia, including 2 to more than 20 nuclei ^6^. Based on these observations, we screened two large libraries of EMA/FDA-approved small molecules to search for drugs that inhibit Spike-induced syncytia formation ^33^. This screening led to the identification of Niclosamide, a synthetic salicylanilide developed in the 1950s as a molluscicide against snails ^34^ and later approved for tapeworm infection in humans, as the most effective drug. The second effective molecule against syncytia formation was Clofazimine, an antibiotic used for the combination treatment of leprosy ^35^ and, more recently, for drug-resistant tuberculosis ^36^.

Of relevance for platelets, we discovered that Niclosamide blocks the formation of syncytia by inhibiting the cellular Ca^2+^-dependent chloride channel and scramblase TMEM16F, thus preventing the externalisation of phosphatidylserine (PS) on the outer leaflet of the cell plasma membrane ^33^. TMEM16F is essential for lipid scrambling in platelets during blood coagulation ^37, 38^. Externalised PS in platelets serves as an anchoring site for the assembly of the tenase and prothrombinase complex, which jointly enhance the rate of thrombin generation by 5 to 10 orders of magnitude ^39^.

Based on these observations, it was tempting to speculate that Spike-driven activation of platelets in SARS-CoV-2 infected lungs could be causally involved in the thrombotic process. Here we show that Spike exposed on the surface of cells or virions induces platelet activation and adhesion. This leads to a pro-coagulant platelet phenotype, characterized by increase in calcium levels, PS exposure and thrombin generation. Both Niclosamide and Clofazimine are remarkably effective at inhibiting Spike-induced platelet activation. Together, these findings provide a molecular mechanism for SARS-CoV-2-induced thrombosis and strengthen the repurposing of these drugs for COVID-19 therapy.

## RESULTS

### SARS-CoV2 Spike potentiates platelet aggregation and adhesion

COVID-19 lung pathology is characterised by extensive thrombosis. Our own analysis of post-mortem samples from over 200 patients who died of COVID-19 at the University Hospital in Trieste, Italy (part of which is reported in ref. ^6^) indicates that thrombosis of the lung microvasculature is present in over 80% of patients with severe COVID-19. Staining of lung sections with an antibody recognising the platelet p62 glycoprotein revealed that these thrombi are massively infiltrated by aggregated platelets (representative images for 4 patients in **Figure 1A).**

**Figure 1.**
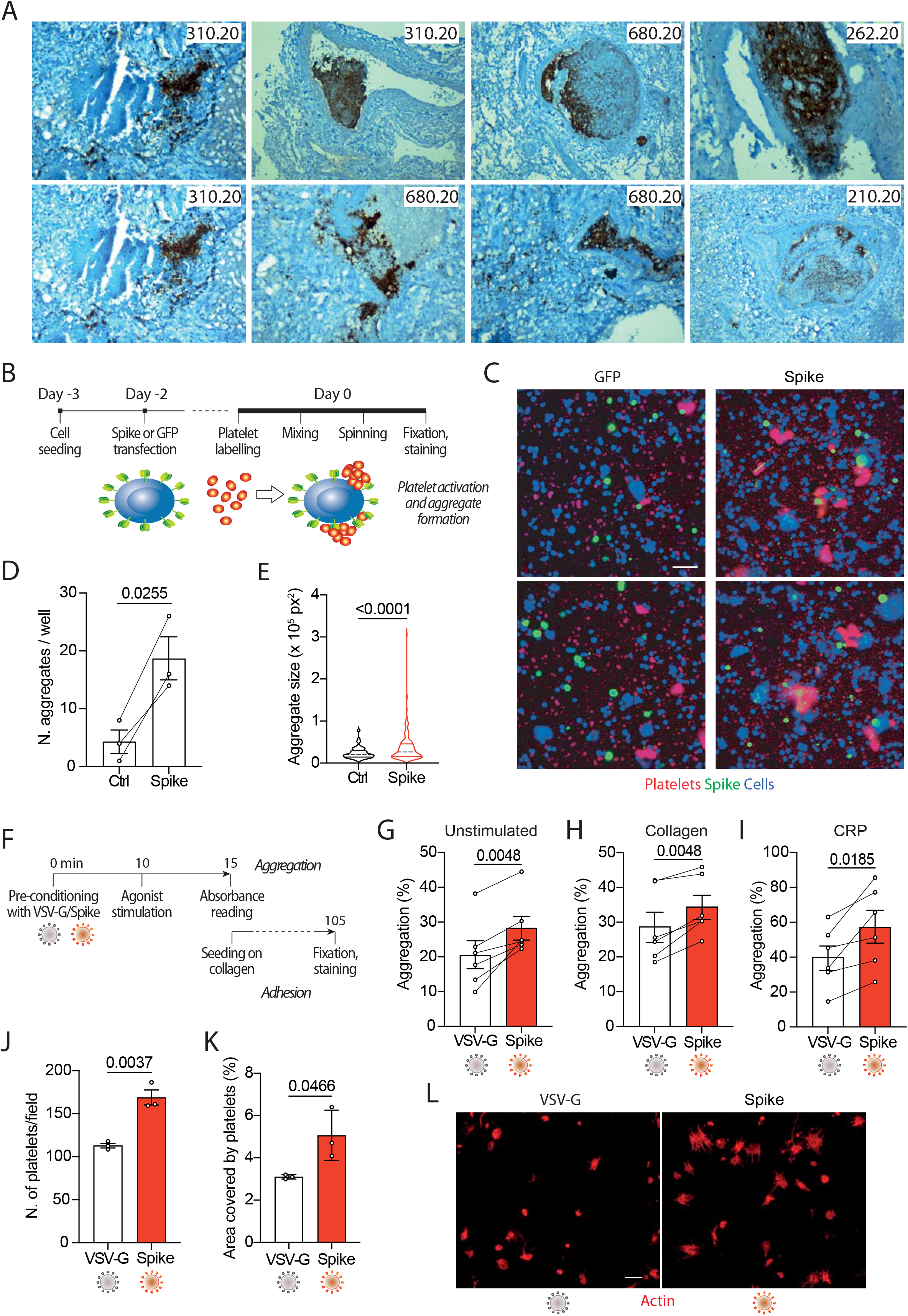
Spike enhances platelet activation. **A.** Histopathological evidence of platelet aggregates in the thrombotic microvasculature of SARS-COV-2-infected lungs from 4 COVID-19 patients. Numeric codes identify patients. Platelets were stained by using an anti p62 glycoprotein antibody. Magnification: x40 **B.** Experimental scheme to study platelet activation and aggregate formation. Vero cells transfected to express either Green Fluorescent Protein (GFP) or SARS-CoV-2 Spike were incubated with pre-labelled washed platelets and shaken at 200 rpm for 10 min at 37°C. The plate was centrifuged, fixed, and stained with Cell Mask and antibodies recognizing either GFP or Spike. **C.** Representative images showing platelet aggregates. Cells stained with Cell Mask are in blue; labelled platelets are in red; GFP or Spike are in green. Scale bar, 20 μm. **D.** Number of aggregates larger than 40,000 px^2^. Results are from n=3 independent experiments. Data are mean ± SEM, statistical significance is indicated (paired Student’s t-test). **E.** Violin plot showing the size of the aggregates found in all the experiments performed. Statistical significance is indicated (unpaired Student’s t-test). **F.** Experimental scheme for platelet activation in suspension. **G-I.** Percentage of platelet aggregation when incubated with vehicle (G), stimulated with collagen (H) or CRP (I). Results are from N=6 independent experiments. Data are mean±SEM Statistical significance is indicated (paired Student’s t-test). **J.** Number of adherent platelets per field. Results are from n=3 independent experiments each performed in duplicate. Each dot represents the mean of 6 images quantified. Data are mean±SEM. Statistical significance is indicated (paired Student’s t-test). **K.** Percentage of the area covered by adherent platelets. Results are from n=3 independent experiments performed in duplicate; each dot represents the mean of 6 images quantified. **L.** Representative images of platelets adhering on collagen. Images were acquired using a high content fluorescent microscope followed by analysis using the ImageJ software (Fiji). Platelets were stained with F-actin (in red). Scale bar, 5 μm.

To understand whether platelet activation might be a direct consequence of the presence of SARS-CoV-2 Spike on the plasma membrane of infected cells, we incubated Vero cells transfected to express Spike (Wuhan strain) or GFP control with washed human platelets (1×10^5^), which had previously been stained with the lipophilic dye CellTracker (scheme in **Fig. 1B).** Transfection efficiency was verified by immunostaining for the transgenes (>40% efficiency in both cases). We observed that the cells expressing Spike significantly increased both the number of platelet aggregates and their overall aggregate area **(Figs. 1D** and **1E** respectively; *P*<0.001 in both cases; representative images in **Fig. 1C).**

Next, we wanted to assess whether Spike was also capable of activating platelets in a cell-free context. We generated lentiviral vectors, containing a GFP-expressing HIV genome, pseudotyped with either SARS-CoV-2 Spike(Δ19) or the vascular stomatitis virus (VSV) G protein as a control. The pseudotyped vector preparations had comparable genome titres **(Suppl. Fig. 1A)** and Spike was detected in the vector lysates by immunoblotting **(Suppl. Fig. 1B).** Both pseudotyped lentiviral preparations were effective at transducing HEK293/ACE2 reporter cells (quantification and representative images in **Suppl. Figs. 1C** and **1D).** As indicators for activation, we measured platelet aggregation and adhesion (scheme in **Fig. 1F).** The Spike pseudoparticles significantly enhanced platelet aggregation (from 21±4% in VSV-G controls to 28±3% in Spike-treated platelets; n=6, *P*<0.01; **Fig. 1G).** There was no effect of VSV-G particle-treated compared to untreated platelets **(Suppl. Fig. 1E** and **1F).** Spike pseudotyped virions also increased aggregation induced by either collagen (29±4% vs. 34±3; n=6, *P*<0.01 in VSV-G vs. Spike; n=6, *P*<0.01) or collagen-related peptide (CRP; 40±7% vs. 57±9%; n=6, *P*<0.01; **Figs. 1H** and **1I** respectively).

In addition to potentiating platelet aggregation, Spike also increased platelet adhesion and spreading. The Spike viral particles augmented both the number of platelets adhering on collagen (113±3 vs. 169±9 platelets per field in VSV-G vs. Spike; n=3, *P*<0.01) and the area covered by the adherent platelets (3% vs. 5±1% in VSV-G vs. Spike; n=3, *P*<0.05); **Figs. 1J** and **1K** respectively; representative images in **Fig. 1L.** Also in this case, there was no significant difference between VSV-G particle-treated and untreated platelets **(Suppl. Fig. 1G).**

Taken together, these observations indicate that exposure of human platelets to SARS-CoV-2 Spike exposed on the surface of either expressing cells or enveloped virions enhances platelet aggregation and adhesion.

### SARS-CoV2 Spike induces a procoagulant response in platelets

Our previous work has shown that expression of Spike leads to activation of members of the TMEM16 family of cellular chloride channels and scramblases ^33^. In particular, cell fusion and formation of large cellular syncytia depends on the Spike-mediated activation of TMEM16F and the consequent exposure of phosphatidylserine (PS) onto the outer leaflet of the plasma membrane. As TMEM16F activation followed by PS exposure is a major mechanism that promotes the procoagulant activity of platelets ^37–39^, we wanted to study PS externalisation in platelets upon treatment with Spike virions (experimental scheme in **Fig. 2A).** We observed that the dual stimulation of platelets with collagen and thrombin in the presence of pseudotyped virions containing Spike significantly increased both the percentage of annexin V-positive platelets (7.0% vs. 10±1% in VSV-G vs. Spike; n=4, *P*<0.05) and the amount of platelet-bound annexin V (1524±98 vs. 1755±82 arbitrary units (AU) in VSV-G vs. Spike; n=4, *P*<0.01; **Figs. 2B** and **2C** respectively; representative flow cytometry profiles are in **Figs. 2D).**

**Figure 2.**
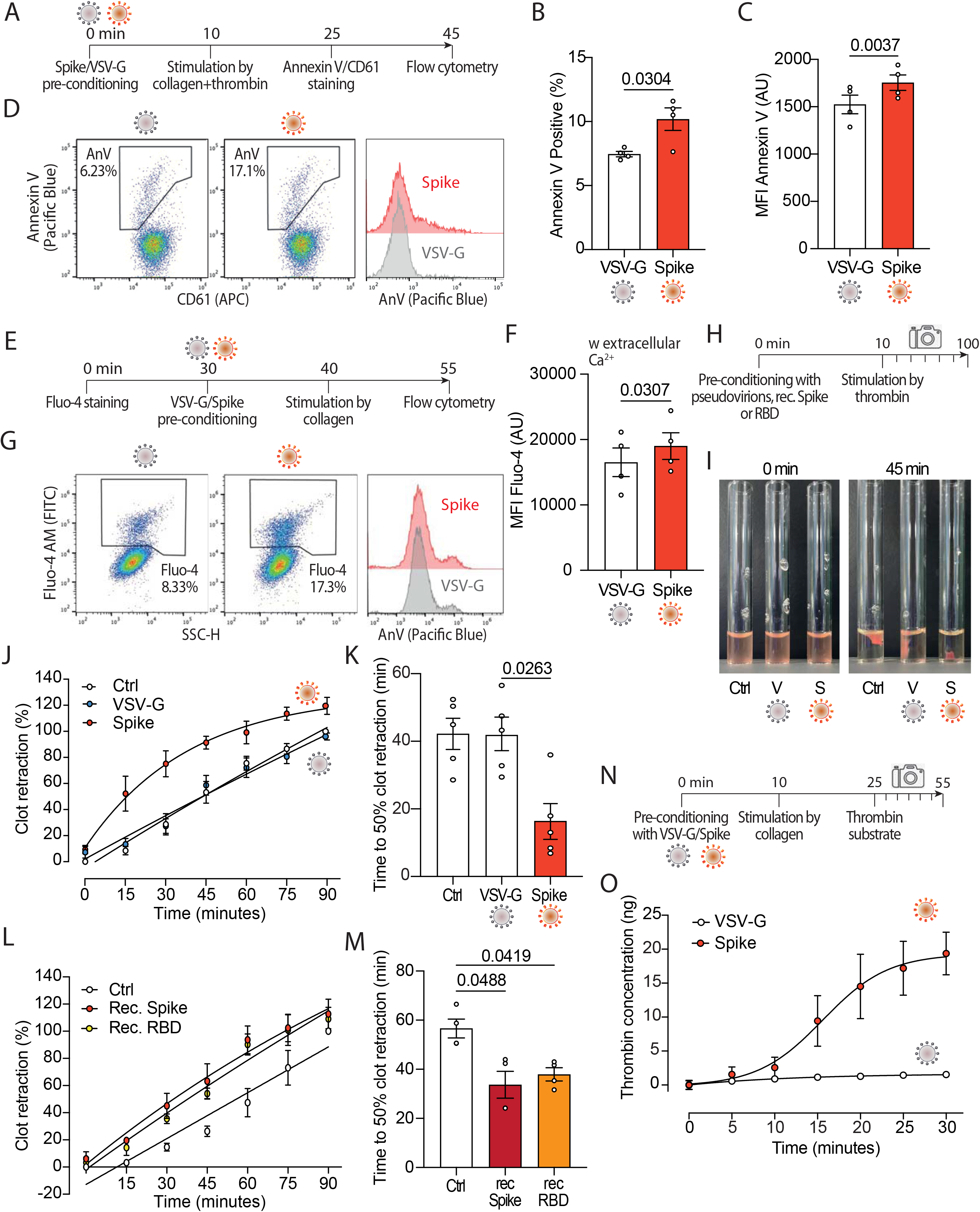
Spikes activates the pro-coagulant activity of platelets. **A.** Experimental scheme to study platelet activation by pseudovirions. Washed platelets were incubated with 1:10 diluted VSV-G or Spike for 10 min, followed by incubation with collagen (30 μg/ml) and thrombin (0.5 units) for 15 min. Platelets were stained with Annexin V-Pacific Blue and CD61-APC for 20 min at 37°C and then analysed by flow cytometry. **B-C.** Percentage and mean fluorescence intensity (MFI) of annexin V positive platelets upon activation with Spike or VSV-G pseudovirions. Results are from n=4 independent experiments. Data are mean±SEM. Statistical significance is indicated (paired Student’s t-test). AU: arbitrary units **D.** Representative flow cytometry plots. The boxed area shows the percentage of platelets positive for annexin V (AnV) binding following incubation with VSV-G or Spike. The histogram on the right shows the distribution of Annexin V positive platelets after the two treatments. **C-D.** Percentage (C) and mean fluorescence intensity (MFI) of annexin V positive platelets upon activation with Spike or VSV-G pseudovirions. Results are from n=4 independent experiments. Data are mean±SEM. Statistical significance is indicated (paired Student’s t-test). AU: arbitrary units **E.** Experimental scheme to study calcium flux in platelets. Washed platelets were stained with Fluo-4 for 30 min, followed by incubation with VSV-G or Spike pseudovirions diluted 1:10 for 10 min. Platelet samples were then incubated with collagen (30 μg/ml) for 15 min and then analysed by flow cytometry. **F.** Mean fluorescence intensity (MFI) of Fluo-4 (AU, arbitrary units) in platelets stimulated with Spike or VSV-G pseudovirions. Results are from n=4 independent experiments. Data are mean±SEM. Statistical significance is indicated (paired Student’s t-test). **G.** Representative flow cytometry plots. The boxed area shows the percentage of platelets positive for Fluo-4 fluorescence following incubation with VSV-G or Spike. The hystogram on the right shows the distribution of Fluo-4 fluorescence after the two treatments. **H.** Experimental scheme for the clot retraction assay following stimulation of platelets with Spike. PRP was supplemented with CaCl_2_ and 10 μl of whole blood, then incubated with 1:10 diluted VSV-G or Spike pseudoparticles, recombinant Spike or recombinant RBD (1 ng/mL) for 10 min at 37°C, followed by incubation with thrombin. Clot retraction was measured over 90 min, taking an image every 15 min. **I-K.** Clot retraction of PRP pre-incubated with PBS, VSV-G or Spike pseudoparticles. Representative images immediately after the addition of thrombin and after 45 min are in I, the percentage of clot retraction over a 90 min observation period is in J. The graph in K shows the time to 50% clot retraction. All data are mean± SEM from n=5 independent experiments. Statistical significance is indicated in K (one-way ANOVA with Dunnett’s post-hoc correction for multiple comparisons). **L-M.** Clot retraction of PRP pre-incubated with PBS, recombinant Spike (1 ng/mL) or recombinant receptor binding domain (RBD, 1 ng/mL). The percentage of clot retraction over the 90 min observation period is in L. The graph in M shows the time to 50% clot retraction in the three experimental conditions. All data are mean± SEM from n=3 independent experiments. Statistical significance is indicated in K (one-way ANOVA with Dunnett’s post-hoc correction for multiple comparisons). **N.** Experimental scheme to study thrombin generation upon platelet treatment with Spike pseudovirions. PRP was incubated with 1:10 diluted VSV-G or Spike pseudovirions for 10 min at 37°C, followed by stimulation with collagen (30 μg/mL) for 15 min (350 rpm, 37°C) and the addition of a fluorogenic thrombin substrate. Thrombin activity was assessed by measuring the conversion of thrombin substrate into its fluorogenic state using a CLARIOstar fluorescent plate reader with 350/450 nm excitation and emission filter. Analysis was performed using MARS analysis software. **O.** Concentration of thrombin formed during a 30 min-time period. Results are from n=5 independent experiments. Data are mean±SEM.

Given the importance of calcium signalling in platelet activation and pro-coagulant activity, we also investigated the effect of Spike on calcium flux (experimental scheme in **Fig. 2E).** We observed that, in platelets activated with collagen, Spike increased the levels of the Fluo-4 indicator, which is sensitive to cytosolic calcium (16,756±2.115 AU vs. 19,281±1952 AU in VSV-G vs. Spike; n=4, *P*<0.05; **Fig. 2F**; representative flow cytometry plots are in **Figs. 2G).** Of interest, this increase was only observed in the presence of extracellular calcium, suggesting that Spike-mediated platelet activation is not related to Ca^2+^ release from the platelet intracellular stores **(Suppl. Fig. 2).**

In light of the effects of Spike on PS exposure and calcium flux, we wanted to assess whether Spike also affected clot formation and retraction, which are the last steps in the coagulation cascade (scheme in **Fig. 2H).** We observed that Spike markedly enhanced thrombin-induced clot retraction (representative images in **Fig. 2I**; complete time point set in **Suppl. Fig. 3).** In particular, the kinetics of clot formation in the presence of Spike was faster compared to both VSV-G-treated or control platelets **(Fig. 2J**; difference in Area Under the Curve, AUC: *P*<0.001 for both controls), with a time to 50% clot retraction of 42±5 min compared to 16±5 min after treatment with VSV-G; n=4, *P*<0.05 **(Fig. 2K).** The same result was also obtained by measuring the clot retraction after incubating platelets with recombinant proteins corresponding to full-length Spike or to the Spike receptor binding domain (RBD) **(Figs. 2L** and **2M** for kinetic analysis and time to 50% clot retraction, respectively; *P*<005 in both cases).

Next, we measured thrombin generation following stimulation with collagen (scheme in **Fig. 2N).** We found that the kinetics of thrombin formation was markedly increased by incubation with the Spike virions (1.5±0.4 ng vs. 19.4±3.1 ng thrombin concentration in VSV-G vs. Spike at 30 min; n=5, AUC difference: *P*<0.01; **Fig. 2O).** There was no significant difference between untreated and VSV-G-treated platelets at any time point (not shown).

Collectively, these results indicate that SARS-CoV-2 Spike markedly increases platelet Ca^2+^ flux, PS exposure and thrombin-induced clot retraction, all of which are markers of the pro-coagulant function in platelets.

### Drugs inhibiting Spike-induced syncytia formation also block the effect of Spike on platelets

Our previous screening work has identified Niclosamide and Clofazimine as the two most potent drugs that inhibit Spike-induced cell fusion by targeting TMEM16F ^33^. As this scramblase is abundant in platelets (immunoblotting from 6 normal donors in **Suppl. Fig. 4A)** and is known to orchestrate the pro-coagulant response ^37–39^, we hypothesized that Niclosamide and Clofazimine may inhibit Spike-induced platelet activation (scheme in **Fig. 3A).**

**Figure 3.**
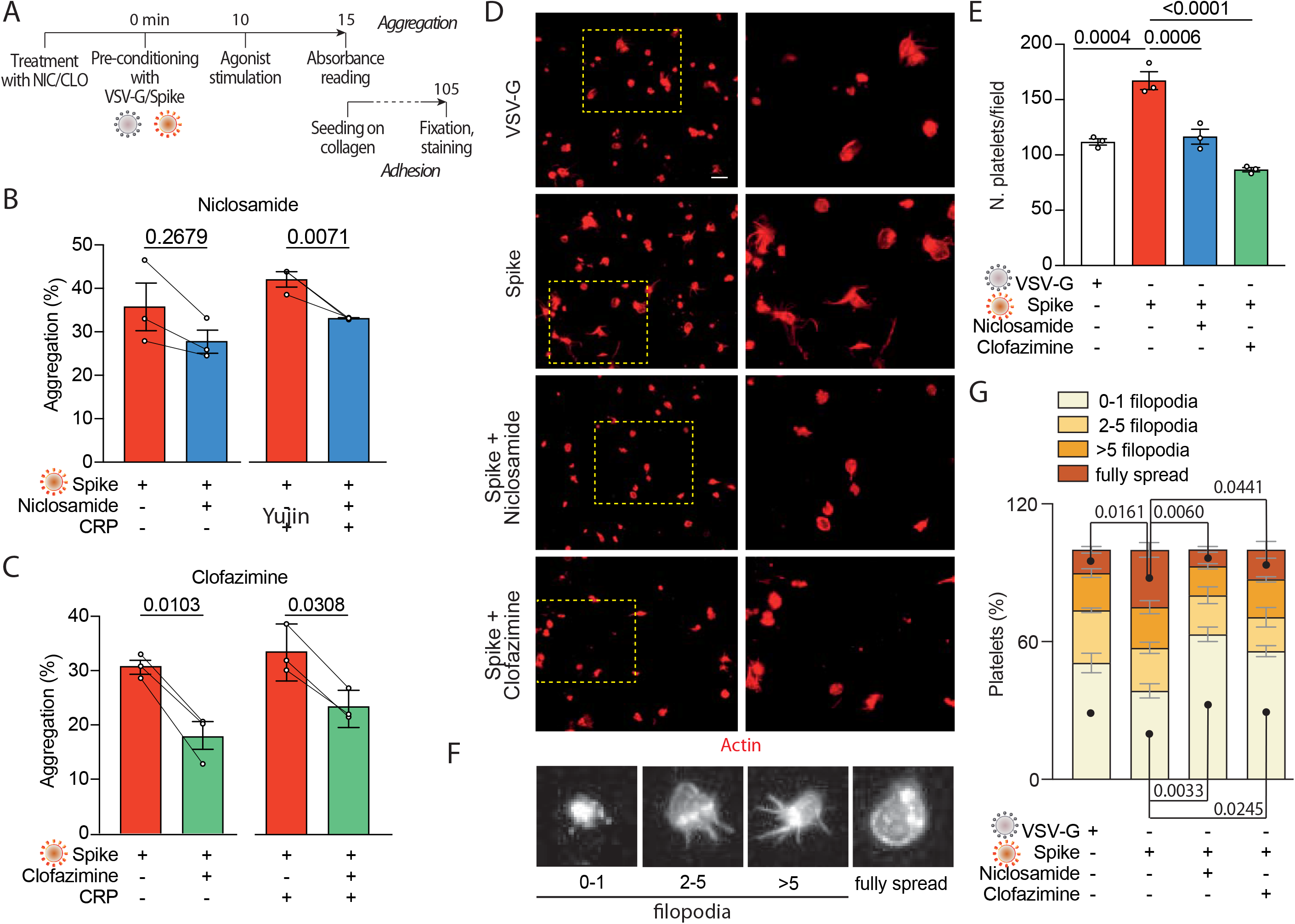
Niclosamide and Clofazimine inhibits spike-induced activation of platelets. **A.** Experimental scheme to assess the effect of drugs on platelet activation. Washed platelets were incubated with Niclosamide, Clofazimine (C) or vehicle for 10 min, then incubated with 1:10 diluted Spike or VSV-G pseudoparticles for further 10 min. Aggregation and adhesion were evaluated as described in Figure 1. **B-C.** Percentage of platelet aggregation upon treatment with Spike in the presence of either Niclosamide (B) or Clofazimine (C), and respective DMSO controls, with or without CRP. Results are from n=3 independent experiments. Data are mean±SEM. Statistical significance is indicated (paired Student’s t-test). **D.** Representative images of platelets adhering on collagen (magnification in the right panels). Platelets are stained in red for F-actin. Scale bar, 5 μm. **E.** Number of adherent platelets per field. Results are from n=3 independent experiments performed in duplicate. Each dot represents the mean of 6 quantified images. Data are mean±SEM. Statistical significance is indicated (unpaired Student’s t-test). **F.** Representative images of platelet morphological changes. Adherent platelets were classified into four morphological categories representing different stages of platelet adhesion and activation: platelets with 0-1, 3-5, more than 5 protrusions or fully spread platelets, as indicated by the representative images under the graph. **G.** Platelet morphological changes upon Spike and drug treatment. Results are from n=3 independent experiments performed in duplicate. Data are mean±SEM. Statistical significance is shown for the indicated morphological categories (one-way ANOVA with Dunnett’s post-hoc correction for multiple comparisons).

Pre-treatment of platelets with either drug, followed by incubation with VSV-G or Spike pseudotyped particles, significantly reduced platelet aggregation upon stimulation with CRP (shown in **Fig. 3B** for Niclosamide - from 42±2% to 33%; *P*<0.05, and in **Fig. 3C** for Clofazimine; from 34±3% to 23±2%; *P*<0.01). Clofazimine also inhibited aggregation in the absence of CRP treatment (from 31±1% to 18±3%; *P*<0.01). There was no effect when the drugs were used in the presence of the VSV-G pseudotypes **(Suppl. Figs. 4B** and **4C).**

We next sought to understand whether Niclosamide and Clofazimine affected Spike-induced platelet adhesion on collagen. Pre-treatment with either drug significantly reduced the total number of adherent platelets (representative images and quantification in **Figs. 3D** and **3E**; 169±9 platelets/field in vehicle-treated samples vs. 118±7 and 88±2 platelets/field in Niclosamide and Clofazimine-treated platelets respectively; n=3, *P*<0.01 in both cases). Pre-treatment with neither Niclosamide nor Clofazimine affected platelet adhesion in VSV-G controls **(Suppl. Fig. 4D).**

Finally, we assessed the spreading stage of adherent platelets as an indicator of drug efficacy. In VSV-G control-treated samples most of the platelets showed a rounded appearance with no or single filopodia (filopodia patterns are in **Fig. 3F).** Platelet activation by the Spike pseudovirions increased the percentage of platelets with multiple filopodia and of platelets showing a fully spread phenotype **(Fig. 3G).** Treatment with either Niclosamide and Clofazimine significantly reversed this Spike-induced phenotype (n=3 per condition; *P*<0.05 for both platelet phenotype distribution between VSV-G and Spike and for the effect of either drug vs. Spike).

### Niclosamide and Clofazimine block Spike-induced platelet procoagulant activity

Given the reduction in aggregation and adhesion, we next investigated whether Niclosamide and Clofazimine could also inhibit the pro-coagulant response induced by Spike (scheme in **Fig. 4A).** Pre-treatment of platelets with either drug caused a significant reduction in PS exposure, as measured by annexin V (AnV) binding (2467±125 AU vs. 1627±55 AU in Spike vs. Niclosamide; 2064±343 AU vs. 1341±113 AU in Spike vs. Clofazimine; n=4, *P*<0.05 in both cases; representative flow cytometry plots and quantification in **Figs. 4B** and **4C).** As increases in intracellular calcium are essential for platelet activation and for TMEM16F activity, we tested whether Niclosamide or Clofazimine could inhibit the increased calcium flux observed following stimulation by Spike and collagen (scheme in **Fig. 4D).** Consistent with the drug effect on platelet activity, both drugs significantly reduced Fluo-4 fluorescence (20,014±2,627 AU vs. 16,607±2,278 AU in Spike vs. Niclosamide; 15,605±2,083 AU vs. 13,804±1,816 AU in Spike vs. Clofazimine; n=4, *P*<0.05 in both cases; representative plots and quantification in **Figs. 4E** and **4F).**

**Figure 4.**
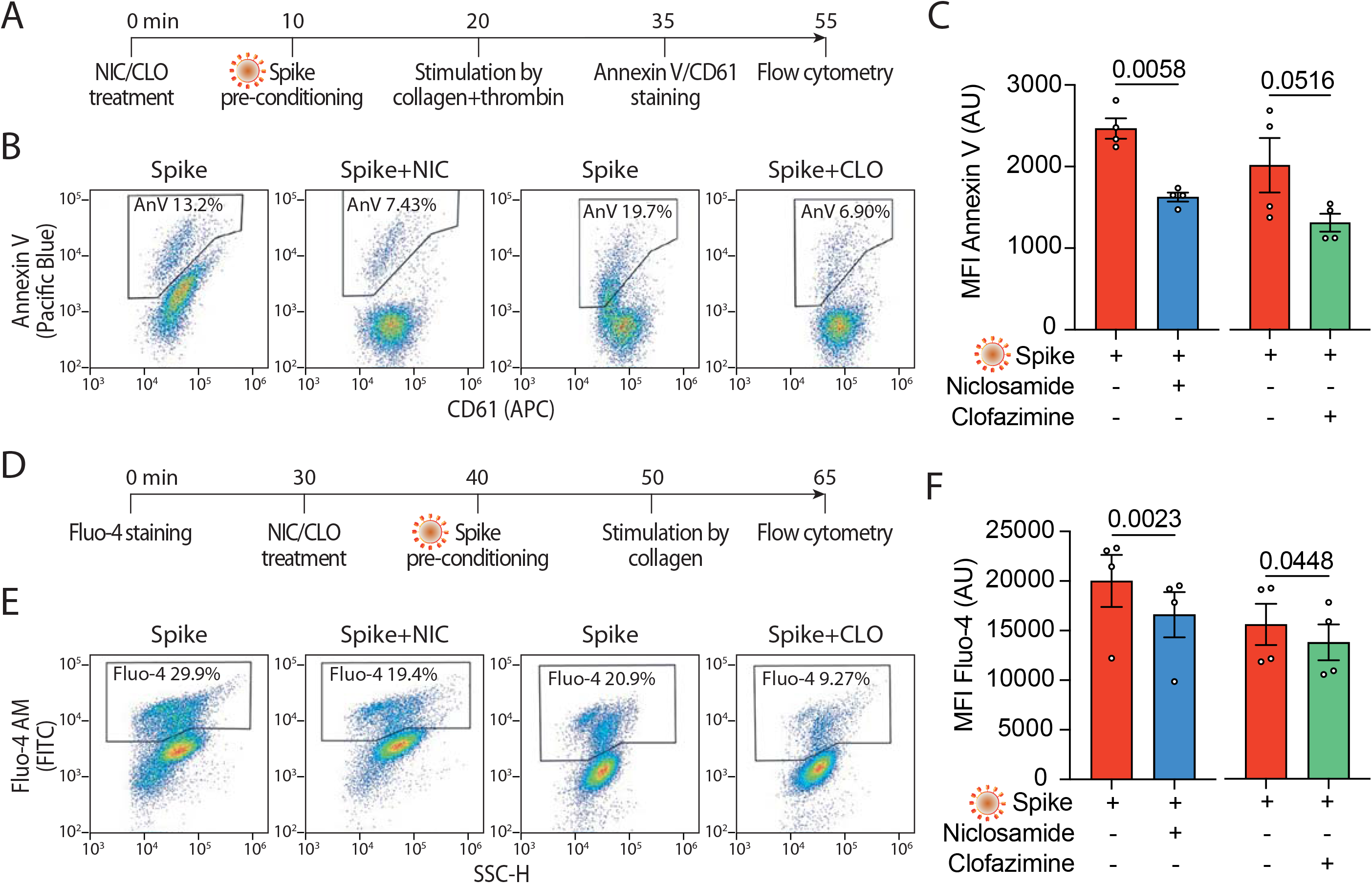
Niclosamide and Clofazimine reduce annexin V and intracellular calcium. **A.** Experimental scheme to assess annexin V reactivity upon Spike stimulation and drug treatment. Platelets were pre-incubated with Niclosamide (NIC, 1 μM) or Clofazamine (CLO, 5 μM) for 10 min, followed by incubation with collagen (30 μg/ml) and thrombin (0.5 units) for 15 min. Platelets were then stained with Annexin V-Pacific Blue and CD61-APC and analysed by flow cytometry. **B.** Representative flow cytometry plots. The boxed areas show the percentage of washed platelets positive for annexin V upon pre-treatment with Niclosamide (NIC), Clofazimine (CLO) or vehicle and incubation with VSV-G or Spike pseudoparticles, followed by stimulation with collagen and thrombin. **C.** Mean fluorescence intensity (MFI) of annexin V positive platelets (AU, arbitrary units). Results are from n=4 independent experiments. Data are mean±SEM. Statistical significance is indicated (paired Student’s t test). **D.** Experimental scheme to assess calcium influx upon Spike stimulation and drug treatment. Platelets were were stained with Fluo-4 for 30 min and then pre-incubated with Niclosamide (NIC, 1 μM) or Clofazimine (CLO, 5 μM) for 10 min, followed by incubation with collagen (30 μg/ml) and thrombin (0.5 units) for 15 min. Platelets were then assessed for fluorescence by flow cytometry. **E.** Flow cytometry plots. The boxed areas show the percentage of washed platelets positive for Fluo-4 upon pre-treatment with Niclosamide (NIC), Clofazimine (CLO) or vehicle (DMSO) and incubation with VSV-G or Spike pseudoparticles, followed by stimulation with collagen and thrombin. **F.** Mean fluorescence intensity (MFI) of Fluo-4 (AU, arbitrary units). Results are from n=4 independent experiments. Data are mean±SEM. Statistical significance is indicated (paired Student’s t test).

Both Niclosamide and Clofazimine caused a significant reduction in the rate of thrombin-stimulated clot retraction (experimental scheme in **Fig. 5A** and representative images in **Figs. 5B** and **5C** for Niclosamide and Clofazimine respectively; complete time courses in **Suppl. Figs. 5A** and **5D).** In the presence of either drug, the kinetics of Spike-induced clot formation was reduced **(Figs. 5D** and **5E** for Niclosamide and Clofazimine respectively; AUC difference: *P*<0.001 and *P*<0.05 for the two drugs respectively). The time to 50% clot retraction increased from 27±4 min to 76±6 min in the presence of Niclosamide and from 27±7 min vs. 72±17 min in the presence of Clofazimine (n=3, *P*<0.05 in both cases; **Figs. 5F** and **5G).**

**Figure 5.**
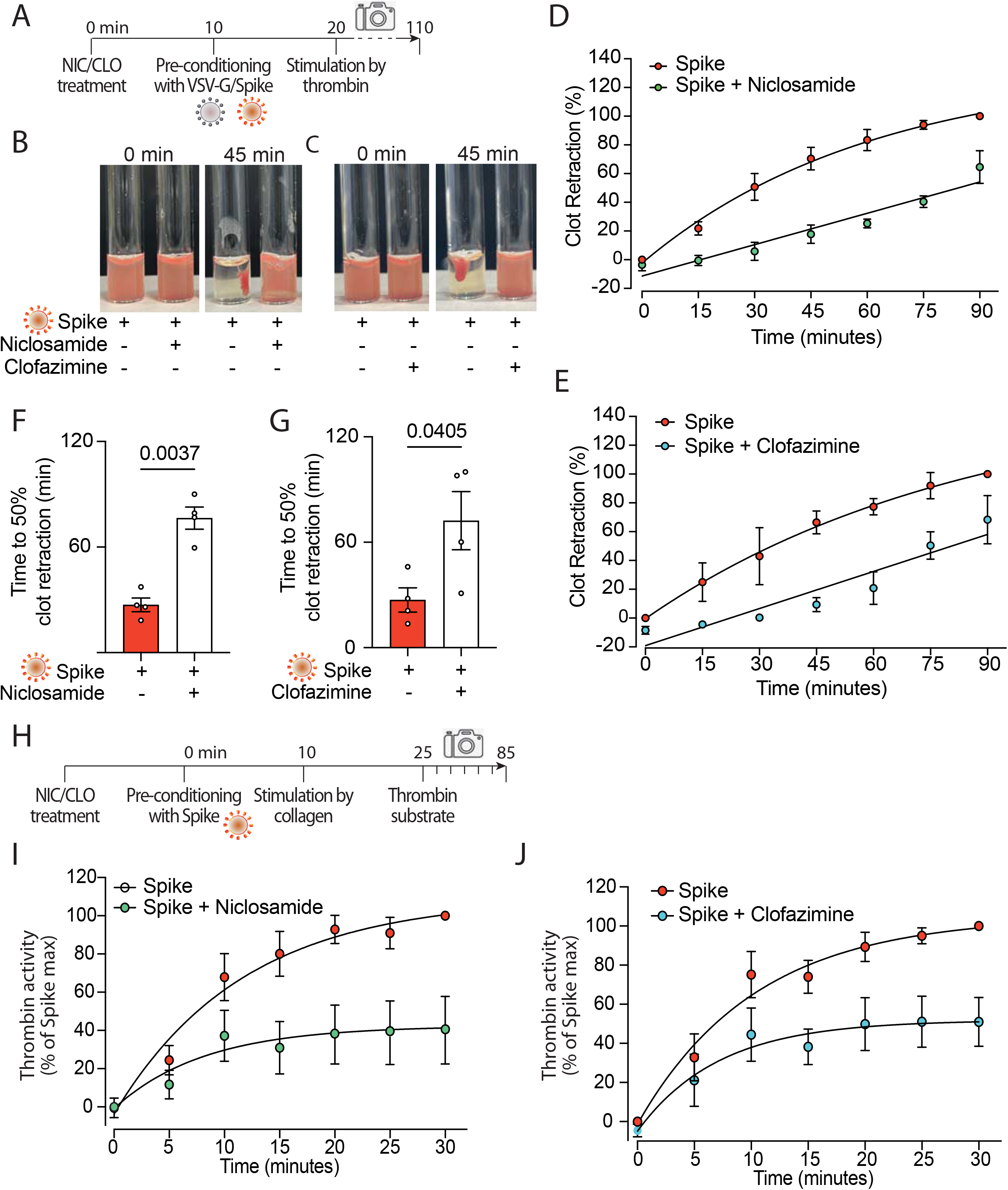
Niclosamide and Clofazimine block pro-coagulant activation of platelets. **A.** Experimental scheme to study the effect of drugs on the clot retraction assay following stimulation of platelets with Spike. PRP, supplemented with CaCl_2_ and 10 μl of whole blood, were incubated with Niclosamide or Clofazimine for 10 min, then treated with 1:10 diluted VSV-G or Spike pseudoparticles for additional 10 min. Clot retraction was measured over 90 min from the addition of thrombin, taking an image every 15 min. **B-C.** Representative images of clot retraction when platelets were treated with Spike and either Niclosamide (B) or Clofazimine (C), immediately after the addition of thrombin and after 45 min. **D-E.** Percentage of clot retraction over 90 min, when platelets were treated with Spike and either Niclosamide (J) or Clofazimine (K). Results are from n=4 independent experiments. Data are expressed as mean ± SEM. **F-G.** Time to 50% clot retraction when platelets were treated with Spike and either Niclosamide (L) or Clofazimine (M). Results are from n=4 independent experiments. Data are mean±SEM. Statistical significance is shown (paired Student’s t-test). **H.** Experimental scheme to study the effect of drugs on thrombin generation upon platelet treatment with Spike pseudovirions. PRP was incubated with 1:10 diluted VSV-G or Spike for 10 min at 37°C, followed by incubation with collagen (30 μg/mL) for 15 min and the addition of a fluorogenic thrombin substrate. Thrombin activity was assessed by the measurement of the conversion of the thrombin substrate into its fluorogenic state. **I-J.** Concentration of thrombin formed during a 30 min-time period in PRP treated with Spike and pre-conditioned or not with Niclosamide or Clofazimine. Data are expressed as mean±SEM.

Finally, we assessed the effect of the two drugs on the kinetics of thrombin generation following stimulation with collagen in the presence of virions with Spike (scheme in **Fig. 5H).** We found that both drugs markedly decreased thrombin formation (62% decrease vs. Spike alone at 30 min for Niclosamide, 55% decrease for Clofazimine; n=5, *P*<0.01 at all time points from 10 to 30 min; **Figs. 5I** and **5J**; AUC difference: *P*<0.01 in both cases).

## DISCUSSION

Here we show that exposure of platelets to the SARS-CoV-2 Spike protein promotes their activation and adhesion, and enhances calcium release and PS exposure to drive increased thrombin generation. SARS-CoV-2 Spike stimulated platelets both when present on the virion envelopes or upon expression onto the plasma membrane of cells. Spike was effective both as a sole agonist or by enhancing the effect of known platelet activators, such as collagen and CRP. Even more notable than the increase in adhesion and aggregation was the effect of Spike on markers of pro-coagulant platelet activation, including PS exposure on the platelet outer membrane and generation of thrombin. A recent report has considered Spike-induced PS externalisation as an indication of platelet apoptosis during COVID-19 ^40^. We instead propose that this is a specific indication of excessive platelet activation, because externalised, negatively charged PS can act as a platform for assembly of the tenase and prothrombinase complex, which massively amplify thrombin generation ^39^.

In SARS-CoV-2 infected patients with severe disease, viral replication is robust in the lung and lower tract respiratory epithelium, leading to the continuous production of infectious particles ^41^. Notably, the lungs of infected individuals contain a significant number of infected pneumocytes and endothelial cells ^42^, which persist for prolonged periods during infection showing expression of viral antigens including Spike ^6^. Cell surface expression of Spike leads to fusion of the infected cells with neighbouring cells expressing the ACE2 receptor, as SARS-CoV-2 Spike, in contrast to the homologous protein from SARS-CoV, contains a furin cleavage site that allows protein activation while the protein is produced during the ER-Golgi transit or when at the cell surface ^43–45^. The large syncytia formed because of these characteristics can be found in over 90% of patients with severe infection and express detectable amounts of Spike on their surface ^33^. We propose that these infected cells and syncytia represent a platform for platelet activation and the consequent induction of thrombosis.

Of note, in our experiments Spike was active both when expressed from cells or on the surface of virions, but also when administered as a recombinant protein. This raises the question as to what the mechanism is for Spike-mediated platelet activation. Based on our previous observations on the activation of TMEM16F by Spike ^33^, we can envisage at least two mechanistic possibilities. First, activation could occur directly upon binding of Spike to the ACE2 receptor, which is expressed in platelets ^40, 46^, leading to direct activation of TMEM16F on the platelet plasma membrane. Second, activation of TMEM16F could be triggered by the increase on Ca^2+^ we observe after stimulation with Spike. In other cell types we have analysed, Spike-mediated TMEM16 activation increases the amplitude of spontaneous Ca^2+^ signals ^33^. This is in line with previous reports showing that both TMEM16A and TMEM16F augment intracellular Ca^2+^ signals by increasing the filling of ER stores and augmenting IP3R-induced Ca^2+^ release ^47^. Two of the experiments that we report here, however, favour the former possibility, namely that Spike-induced platelet activation is consequent to a molecular event occurring at the plasma membrane. First, activation of platelets did not occur when the medium was depleted of extracellular Ca^2+^, likely indicating that intracellular Ca^2+^ stores are not required for activation. Second, and most important, we observed that platelet activation also occurred upon treatment with the isolated spike RBD domain. These observations are in favour of a direct TMEM16F activation event occurring upon binding of Spike to the ACE2 receptor. The nature of the relationship and the possible direct interaction between ACE2 and TMEM16F are currently one of the subjects of our investigation.

Both Niclosamide and Clofazimine, which block Spike-induced syncytia formation in a variety of epithelial and non-epithelial cells expressing the ACE2 receptor, were also remarkably effective at inhibiting Spike-induced platelet activation. This is in line with the known role of TMEM16F in amplifying the rate of platelet pro-coagulant activity ^39^ and is consistent with the effect we observed for the two drugs on calcium flux, PS exposure and thrombin generation. Niclosamide was particularly effective at inhibiting platelet activation, acting at concentrations in the low hundred nanomolar range which are even lower than those needed to inhibit syncytia ^33^.

Niclosamide is a synthetic salicylanilide developed in the 1950s as a molluscicide against snails ^34^ and subsequently approved for use in humans, where it has been employed for over 50 years to treat tapeworm infections ^48^. Solubility of the currently available oral formulation is relatively low, but there is anyhow evidence of significant systemic absorption, with plasma levels that can reach 1-20 μM ^49^. In a colorectal cancer clinical trial ^50^, administration of 2 g daily Niclosamide per day resulted in plasma concentrations of ^~^0.5-1 μg/ml (https://ascopubs.org/doi/abs/10.1200/JCO.2018.36.15_suppl.e14536), corresponding to ^~^1.5-3 μM, which is far above the concentration we found to inhibit platelet activation by Spike.

Finally, an important question that still remains open is whether the mechanism for Spike-induced platelet activation could be causally linked to the conditions of thrombocytopenia that are commonly observed in COVID-19 patients, even in the absence of signs of consumption ^5, 26–28^. Decreased platelet levels due to clearance in the spleen are reported in other conditions of PS overexposure ^51^, including in mice with reduced levels of Bcl-xL (an anti-apoptosis Bcl-2 family protein member) ^51^. Recent evidence links this feature specifically to TMEM16F and extracellular Ca^2+^ influx. Mice with sphingomyelin synthase 1 deficiency show marked thrombocytopenia due to increased PS exposure consequent to excessive TMEM16F activation ^52^. Based on these observations, we speculate that thrombocytopenia in the course of severe COVID-19 may be a consequence of abnormal platelet activation through Spike-mediated TMEM16F stimulation.

In conclusion, the findings we describe here provide a pathogenic mechanism to explain thrombosis associated to SARS-CoV-2 lung infection and strengthen the proposal of repurposing Niclosamide for COVID-19 therapy.

## ACKNOWLEDGMENTS

This work was supported by grants from the King’s College London King’s Together programme and National Institute for Health Research Biomedical Research Centre at Guy’s & St Thomas’ NHS Foundation Trust and King’s College London for COVID-19 research, with the contribution of the British Heart Foundation (BHF) Programme Grant RG/19/11/34633 and the European Research Council (ERC) Advanced Grant 787971 “CuRE”. HEA, MC and AC were supported by grants from the British Heart Foundation (RG/19/8/34500, PG/17/40/33028 and RG/19/11/34633).

## METHODS

### Human blood collection and isolation of platelets

Platelet studies using human samples were conducted according to the principles of the Declaration of Helsinki and approved by St Thomas’s Hospital, London, UK Research Ethics Committee (Ref. 07/Q0702/24). All volunteers were screened prior to entering the study and gave written informed consent. Blood was collected by venepuncture into tri-sodium citrate (3.2%; Sigma) from healthy volunteers (aged 25-40), who had abstained from non-steroidal anti-inflammatory drug consumption for the preceding 14 days. Platelet rich plasma (PRP) was obtained by centrifugation of whole blood (175×g, 15 min, 25°C). Further purification of platelets was achieved by centrifugation of PRP at 1000×g for 10 min at 25°C in the presence of apyrase (0.02 U/mL, Sigma) and prostacyclin (PGI_2_; 2 μM, Tocris) followed by resuspension of the platelet pellet in modified Tyrode’s HEPES buffer (134 mmol/L NaCl, 2.9 mM KCI, 0.34 mmol/L Na_2_HPO_4_, 12 mmol/L NaHCO_3_, 20 mmol/L HEPES and 1 mmol/L MgCI_2_; pH 7.4; Sigma) with glucose (0.1% w/v; Sigma) and bovine serum albumin (BSA, Sigma). Washed platelets were adjusted to a concentration of 3×10^8^/ml, allowed to rest of 30 min and then supplemented with calcium chloride (CaCl_2_; 2 mM, Sigma).

### Patients

Patients’ lung samples are from post-mortem analysis of 4 patients who died of COVID-19 at the University Hospital in Trieste, Italy, after intensive care support. These are from a more extensive cohort of 41 consecutive patients, who all died of COVID-19. Of these, 25 were male and 16 were female; the average age was 77 for men and 84 for women. During hospitalization, all patients scored positive for SARS-CoV-2 on nasopharyngeal swab and presented symptoms and imaging data indicative of interstitial pneumonia related to COVID-19 disease. Extensive characterization of these patients has previously been reported ^6^. Thrombosis of the lung micro- and macro-vasculature was detected in 29/41 (71%) of all the analysed patients and in 83% of patients in intensive care units. Use of these post-mortem samples for investigation was approved by the competent Joint Ethical Committee of the Regione Friuli Venezia Giulia, Italy (re. 0019072/P/GEN/ARCS).

### Pseudotyped viral particles

For Spike pseudotyped lentiviral particle production, we generated the expression plasmid pEC120-S-D19-V5, in which the 19aa amino acids at the Spike C terminus, acting as an ER-retention signal^53^, are replaced by a V5 epitope tag. The modified DNA segment was obtained by recombinant PCR using primers Fw 5’ ACGCGTCGACTTTTGTGGCAAAGGTT, Re 5’ CCCAAGCTTGGGACGCGTCGTTACGTAGAATCGAGACCGAGGAGAGGGTTAGGGATAGGCTTACCACCACCTCCACCGCAGCATGATCCGCATGAGC and cloned into the pEC117-Spike-V5 vector^33^. The construct was verified by Sanger sequencing. Pseudotyped viral particles were produced as follows: HEK293T cell (2.5 × 10^6^) were seeded in a 100 mm-dish and co-transfected by calcium phosphate with 10 μg pLVTHM/GFP (Addgene 12247), 7.5 μg psPAX2 (Addgene 12260), and 6 μg pMD2.G (Addgene 12259) or pEC120-S-D19-V5, for a total of 13.5 μg DNA for each dish. The culture medium containing pseudotyped particles was collected after 48 hr and concentrated 3 times using Vivaspin columns with 100 kDa cutoff (GE Healthcare, #28932363) following the manufacturer’s instructions. Aliquots were stored at-80°C.

For pseudoparticle titration, RNA from pseudoparticle preparations was isolated using the Viral RNA isolation Kit (Takara) following the manufacturer’s instructions. Viral RNA genome content was quantified using the Lenti-X qRT-PCR Titration Kit (Takara) and the Quant-X One-Step qRT-PCR TB Green Kit (Takara). A Lenti-X RNA Control Template was provided by the kit and served to build a standard curve of viral genome copies. Average Ct values from replicates were plotted vs. the copy number in a log scale and the number of copies per ml was calculated as indicated by the manufacturer.

To assess pseudoparticle infectivity, HEK-293T cells were bulk transfected with a plasmid expressing human ACE2 and then seeded in a 96-well plate (3×10^3^ cells per well). The next day 5 μL VSV-G or 10 μL Spike pseudoparticles were added to each well. The plate was fixed at 48 hours, stained with Hoechst (Invitrogen) and imaged by using Perkin Elmer Operetta CLS High Content Fluorescent microscope. Analysis was performed using the ImageJ software (Fiji).

### Platelet aggregation with pseudotyped viral particles and cells

Platelet aggregation was performed as previously published ^54^ with minor modifications. Briefly, washed platelets were incubated with VSV-G or Spike pseudoparticles (1:10) for 10 min followed by stimulation with collagen related peptide (CRP; 0.3 μg/mL) or collagen (0.3 μg/mL, Takeda) or vehicle. Plates were mixed for 5 minutes (1200 rpm, 37°C; BioShake IQ, Q Instruments), and the absorbance was measured at 595 nm using an absorbance microplate reader (Sunrise, Tecan). Aggregation was calculated as percentage change in absorbance.

For platelet aggregation on Spike-expressing cells, Vero cells were kept in culture in Dulbecco’s Modified Eagle Medium (DMEM) supplemented with 10% of foetal bovine serum (FBS) and transfected using FuGENE HD (Promega) transfectant reagent as suggested by the manufacturer either with a plasmid encoding the Green Fluorescent Protein (GFP; pZac-GFP) or a plasmid encoding the full-length SARS-CoV-2 Spike protein with V5 tag (pSARS-COV-2-S). After detaching, 20,000 cells were co-incubated with 5 × 10^5^ pre-labelled platelets (1:2000 Cell Tracker Deep Red Dye, Thermo Fisher Scientific) in each well of a 96-well plate, and shaken at 200 rpm for 10 minutes at 37°C. The plate was centrifuged at 300 g for 10 min and fixed by directly adding paraformaldehyde (PFA, VWR) to the supernatant to a final concentration of 4%. After washing 3 times with PBS, the plate was permeabilized with 0.5% Triton (Sigma) for 10 min, blocked with 1% BSA for 1 hr at room temperature (RT) and stained with anti-GFP antibody (1:1000; Abeam #ab6556) or anti-V5 tag antibody (1:500; Invitrogen #37-7500) for 2 hr at RT, and Cell Mask (1:2000; Thermo Fisher Scientific) for 30 min at RT. 20X magnification images were acquired using a Perkin Elmer Operetta CLS High Content Fluorescent microscope. Analysis was performed using the ImageJ software (Fiji).

### Platelet adhesion and spreading

Washed platelets (3 × 10^8^/mL) were allowed to adhere to type I Horm collagen (100 μg/mL; Takeda) coated plates, for 90 min at 37°C. Samples were fixed with 0.2% PFA for 10 min followed by permeabilization with 0.2% Triton (Sigma) for 5 minutes. Samples were washed with filtered phosphate buffered saline (PBS) and stained with Phalloidin 488 (1:2000; Thermo Fisher Scientific). 63X magnification images were acquired using a Perkin Elmer Operetta CLS High Content Fluorescent microscope. Analysis was performed using ImageJ software (Fiji).

### Flow cytometry analysis of phosphatidylserine exposure

Phosphatidylserine exposure was analysed by assessment of annexin V binding. Washed platelets were incubated with VSV-G or Spike pseudoparticles (1:10) for 10 min followed by activation with collagen (30 μg/ml) and thrombin (0.5 units, Sigma) or vehicle for 15 min (350 rpm, 37°C; BioShake IQ). Platelets were stained with Annexin V-Pacific Blue (1:50; Biolegend) and CD61-APC (1:100; Biolegend) in the presence of annexin binding buffer for 20 min (350 rpm, 37°C; BioShake IQ). Samples were diluted with modified Tyrode’s HEPES buffer and analyzed on the ACEA Novocyte 3000 (ACEA Biosciences Inc.). Analysis was performed using the FlowJo software v.10 (TreeStar Inc).

### Flow cytometry assessment of calcium flux

To analyse calcium flux, washed platelets were stained with Fluo-4 AM (5 μM, Thermo Fisher Scientific) for 30 min at 37°C) followed by incubation with VSV-G or Spike pseudoparticles (1:10) for 10 min. Platelet samples were activated with vehicle or collagen (30 μg/ml) for 15 min. Samples were diluted with modified Tyrode’s HEPES buffer and analyzed on the ACEA Novocyte 3000 (ACEA Biosciences Inc.). Analysis was performed using the FlowJo software v.10 (TreeStar Inc).

### Thrombin measurement

Thrombin activity was measured using a Thrombin Activity Assay Kit (Abcam) as per the manufacturer’s guidelines. Briefly, PRP was incubated with VSV-G or Spike pseudoparticles (1:10) for 10 min at 37°C, followed by stimulation with collagen (30 μg/ml) for 20 min (350 rpm, 37°C). PRP was diluted 1:10 with thrombin assay buffer, and 50 μl of thrombin substrate was added to each well. Thrombin activity was assessed by measuring the conversion of thrombin substrate into its fluorogenic state using a CLARIOstar fluorescent plate reader (BMG Labtech) with 350/450 nm excitation and emission filter. Analysis was performed using MARS analysis software.

### Clot retraction

PRP was diluted 1:1 with modified Tyrode’s HEPES buffer, supplemented with CaCl_2_ (2 mM) and 10 μl of whole blood. PRP was incubated with VSV-G (1:10), Spike pseudoparticles (1:10), His-tag recombinant SARS-CoV-2 Spike Protein S1/S2 (S-ECD) (1 ng/ml; ThermoFisher Scientific; aa11-1208; RP-87680) or Flag-tag recombinant SARS-CoV-2 Spike RBD (1 ng/ml; Bio-Techne, 10689-CV-100) for 10 min at 37°C, followed by stimulation with thrombin (0.5 units). Clot retraction was measured over 90 min, taking an image every 15 min. Analysis was performed using ImageJ (NIH).

### Effect of drugs on Spike-induced platelet function

Platelet function experiments described above were performed with pre-incubation of platelets with Niclosamide (1 μM) or Clofazimine (5 μM) for 10 min (350 rpm, 37°C). Both drugs were purchased from Sigma. Similar dilutions of DMSO (vehicle) were used as controls.

### Immunoblotting

To detect Spike in pseudoparticles by immunoblotting, concentrated Spike and VSV-G pseudoparticles (40 μL) were diluted in 4x protein loading dye. To visualise TMEM16F, platelets (9×10^8^/ml) were lysed in 2% SDS, quantified by using the Pierce™ BCA Protein Assay Kit (#23225, Thermo Fisher Scientific) and 15-20 μg were resolved by electrophoresis in 4-20% gradient polyacrylamide gels (Mini-PROTEAN, Bio-Rad) and transferred to Trans-blot (Bio-Rad). Membranes were blocked in TBST (TBS ± 0.1% Tween-20) with 5% skim milk (Cell Signaling, 9999) at room temperature for 1 hr. Blots were then incubated (4°C overnight) with primary antibodies against Spike, tubulin or TMEM16F. Blots were washed three times with TBST. Membranes were incubated for 1 hr at room temperature with anti-rabbit HRP-conjugated antibody (1:5,000) or anti-mouse HRP-conjugated antibody (1:10,000). After washing three times with TBST (10 min each), blots were developed with ECL (Amersham).

### Antibodies

**I**mmunofluorescence analysis was performed for actin (AlexaFluor 488 Phalloidin, A12379, ThermoFisher Scientific) and GFP (ab6556, Abcam). Flow cytometry was performed for annexin V (Annexin V-Pacific Blue, Biolegend) and calcium flux (Fluo-4 AM, ThermoFisher Scientific). Immunoblots were performed with primary antibodies against Spike (Genetex, #GTX632604; 1:1,000), tubulin (Cell Signaling, #3873S; 1:10,000) and TMEM16F (Sigma-Aldrich, #HPA038958; 1:1,000).

## SUPPLEMENTARY FIGURE LEGENDS

**Supplementary Figure 1.**
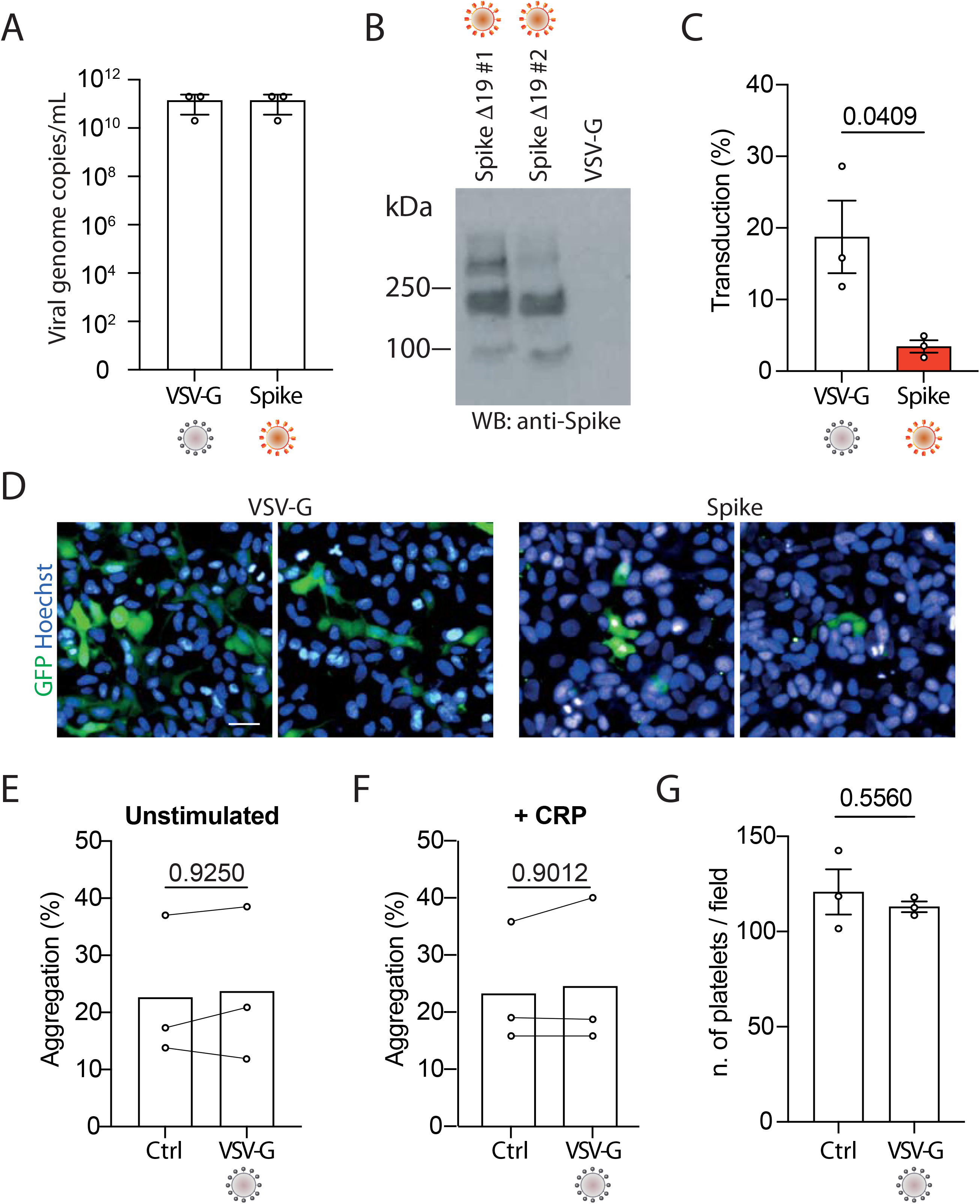
Characterisation and effect of lentiviral vectors pseudotyped with SARS-CoV-2 Spike or VSV-G. **A.** Viral genome titration of pseudoparticles. Results are from the quantifications of n=3 pseudoparticle preparations. **B.** Immunoblot showing the expression of Spike in pseudoparticles. Blots were incubated (4°C, overnight) with a primary antibody recognizing Spike (Genetex, 1:1,000) followed by incubation with an anti-mouse HRP-conjugated antibody (1:10,000) and subsequent development with ECL (Amersham). **C-D.** HEK-293T cells overexpressing the human ACE2 were treated with VSV-G or Spike pseudoparticles and fixed after 48h. Percentage of transduced cells (C) and representative images (D). In green, GFP; in blue, Hoechst. Scale bar, 20 μm. SARS-CoV-2 Spike pseudotyped vectors are known to have a lower efficiency compared to VSV-G-pseudotyped vectors, despite the use of the Δ19 C-terminal deletion, which increases transport of Spike to the plasma membrane ^53^. **E-F.** Platelets were incubated either with VSV-G or PBS for 10 min at 200 rpm, at 37°C and then treated with PBS (E; unstimulated) or stimulated with CRP (F; final concentration 0.3 μg/mL). Aggregation was measured as described in Figure 1. Results are from n=3 independent experiments. Data are mean±SEM. Statistical significance is shown (paired Student’s t-test). **G.** Platelet adhesion was determined as described in Figure 1. Results are from n=3 independent experiments. Data are mean±SEM. Statistical significance is shown (paired Student’s t-test).

**Supplementary Figure 2.**
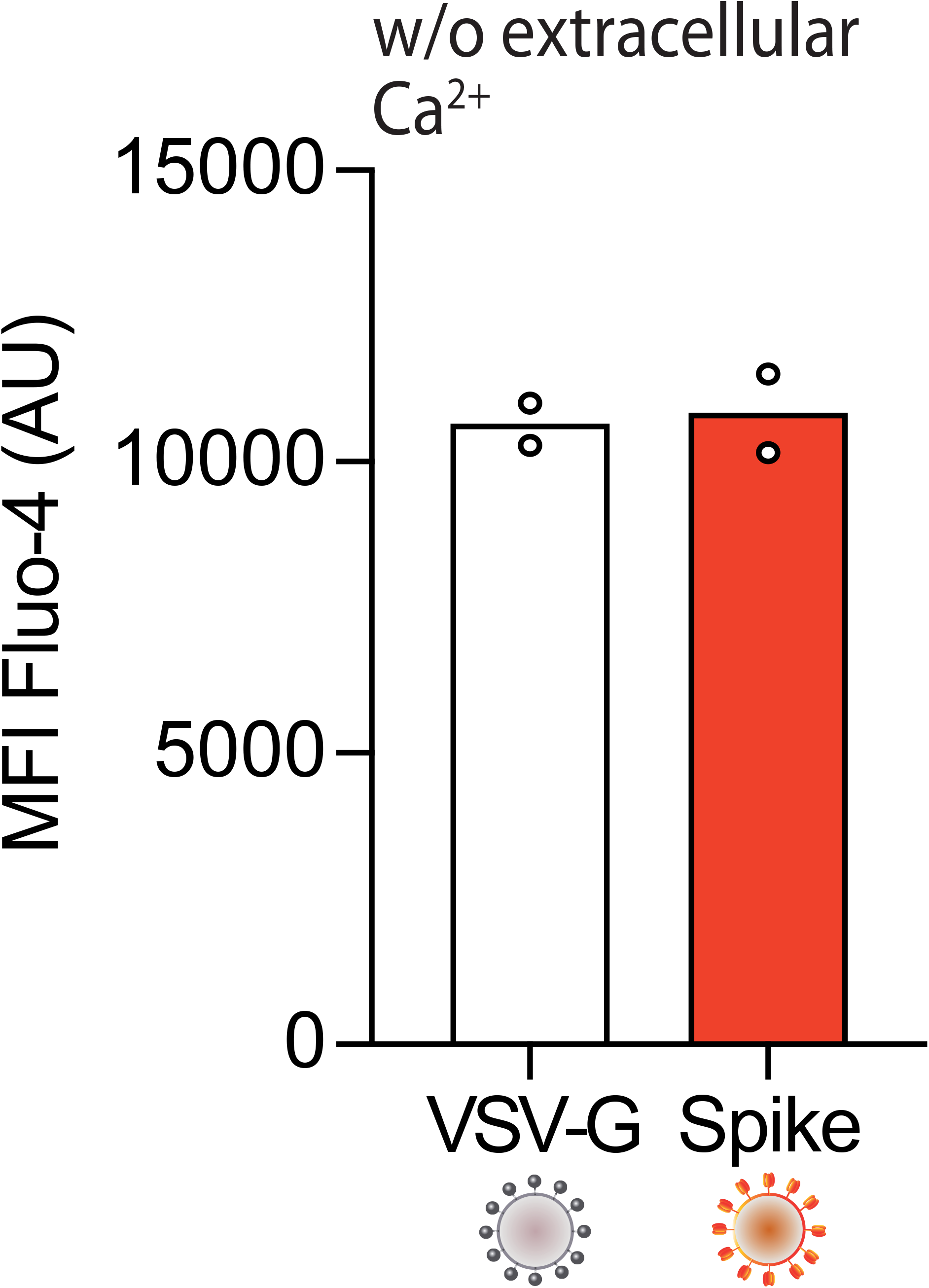
Calcium influx in the absence of extracellular calcium. Washed platelets were stained with Fluo-4 for 30 min, followed by incubation with 1:10 diluted VSV-G or Spike pseudoparticles for additional 10 min. Platelet samples were activated with collagen (30 μg/ml) for 15 min and then analysed by flow cytometry. The graph shows the mean fluorescence intensity (MFI) of Fluo-4 (AU, arbitrary units). Results are from 2 independent experiments. Data are means.

**Supplementary Figure 3.**
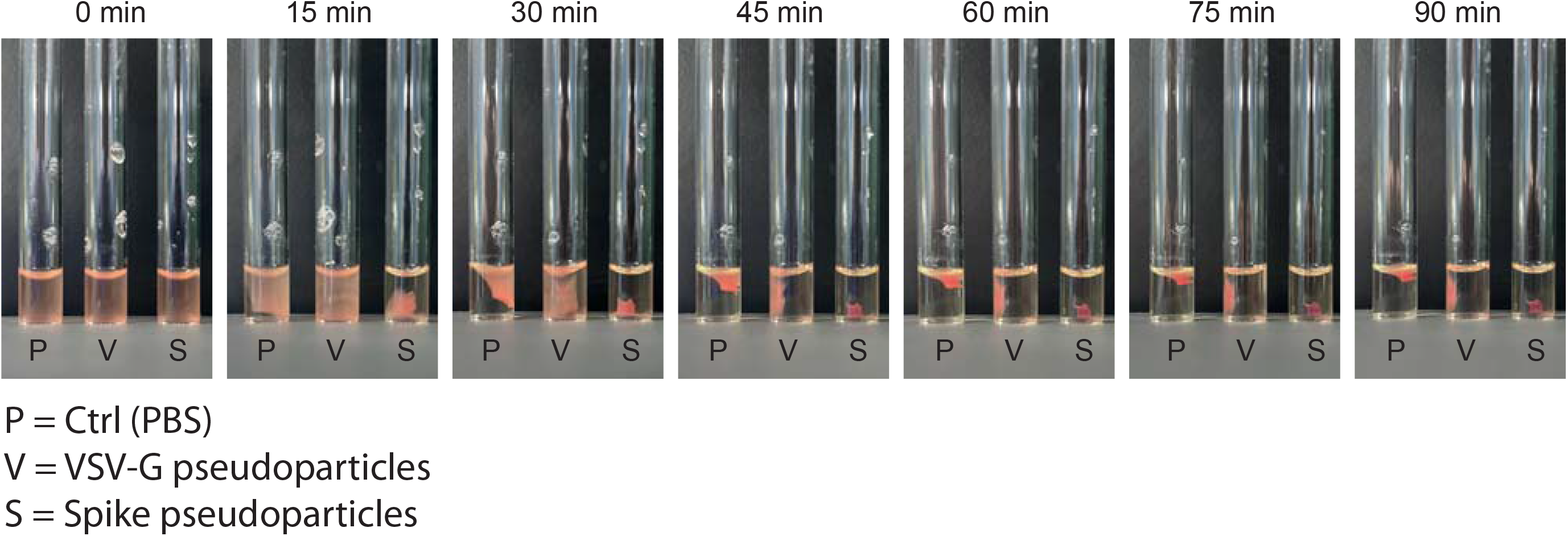
Clot retraction assay. Images of a representative clot retraction assay using PRP incubated with Niclosamide or Clofazimine for 10 min, followed by treatment with 1:10 diluted VSV-G or Spike pseudovirions for additional 10 min. The images were taken every 15 min. P, PBS; V, VSV-G; S, Spike.

**Supplementary Figure 4.**
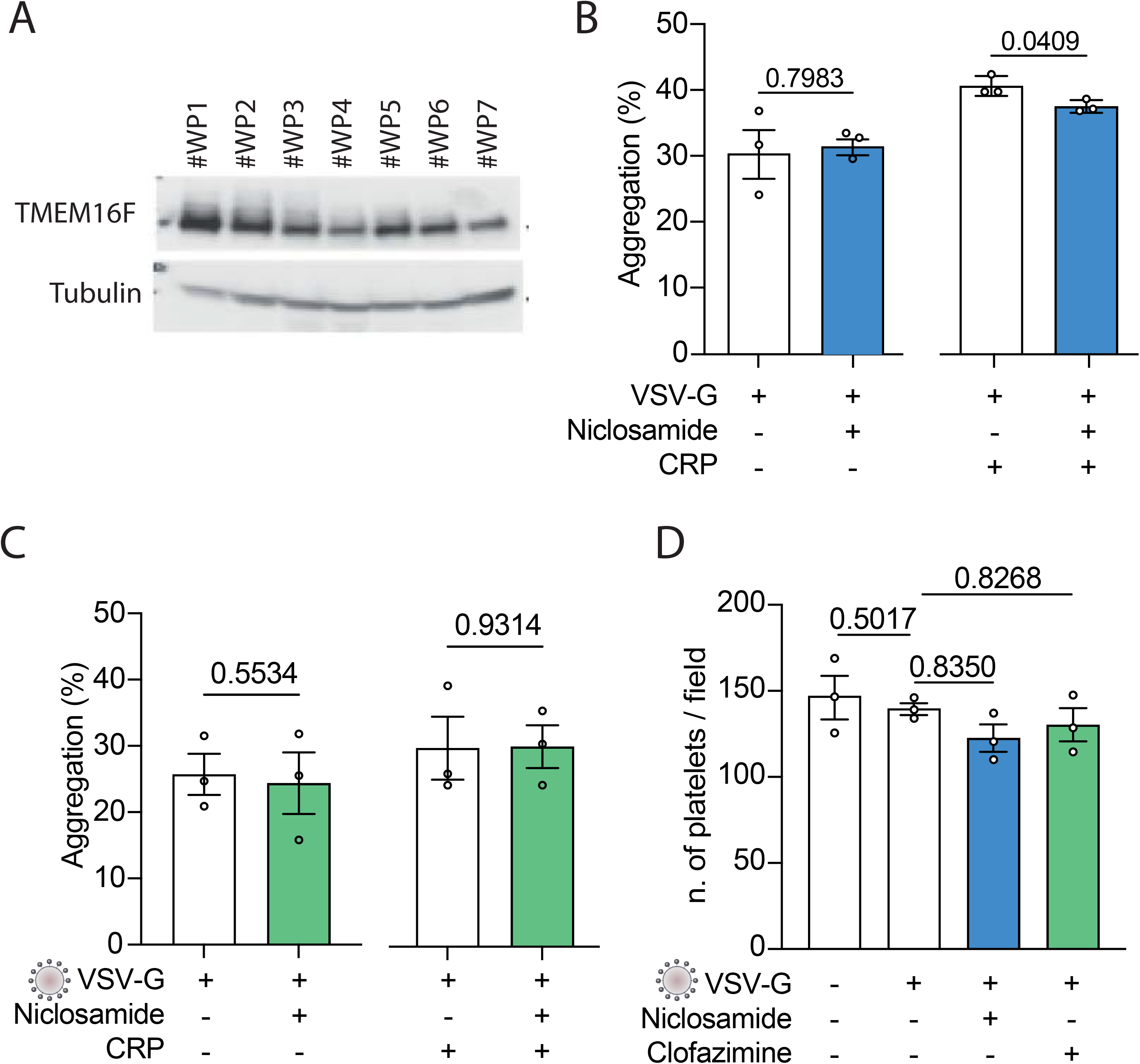
Expression of TMEM16F in platelets and effect of drugs and VSV-G pseudovirions. **A.** Immunoblot showing the expression of TMEM16F by washed platelets (WP) from different normal individuals. Blots were then incubated (4°C, overnight) with primary antibodies recognizing TMEM16F (1:1,000) and tubulin (1:10,000), followed by incubation for 1 hr with either anti-rabbit HRP-conjugated antibody (1:5,000) or anti-mouse HRP-conjugated antibody (1:10,000). ECL (Amersham) was used for blot development. **B.** Platelets were incubated either with Niclosamide or PBS for 10 min, then treated with either VSV-G pseudoparticles or PBS for additional 10 min. Aggregation was measured as described in Figure 3. Results are from n=3 independent experiments. Data are mean±SEM. Statistical significance is shown (paired Student’s t-test). **C.** Platelets were incubated either with Clofazimine or PBS for 10 min, then treated with either VSV-G pseudoparticles or PBS for additional 10 min. Aggregation was measured as described in Figure 3. Results are from n=3 independent experiments. Data are mean±SEM. Statistical significance is shown (paired Student’s t-test). **D.** Platelet adhesion was evaluated as described in Figure 3. Results are from n=3 independent experiments performed in duplicate; 6 images per well were analysed. Data are mean±SEM. Statistical significance is shown (one-way ANOVA with Dunnett’s multiple comparison test).

**Supplementary Figure 5.**
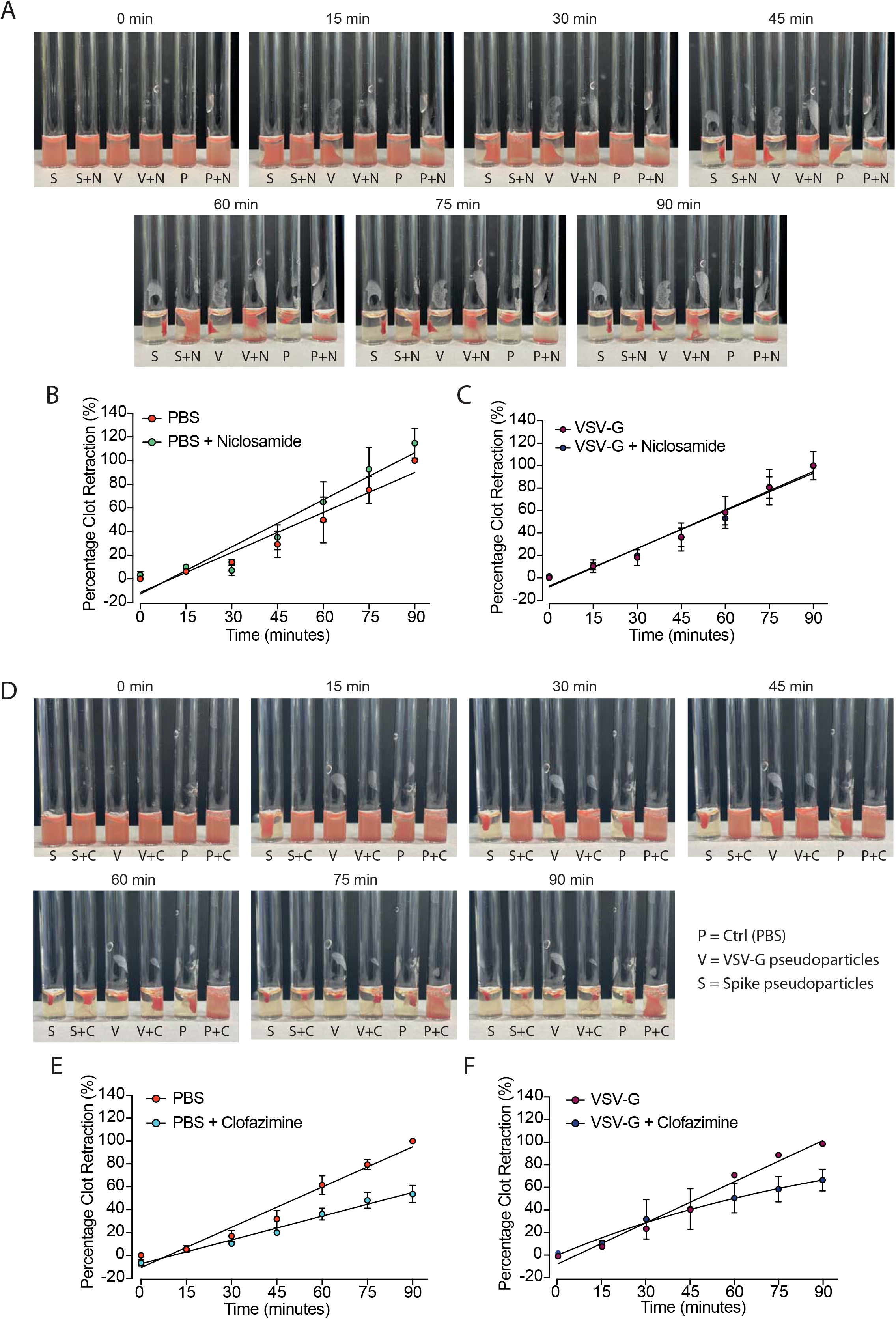
Effect of PBS and VSV-G Controls on clot retraction. **A.** Images of a representative clot retraction assays using PRP incubated with Niclosamide or Clofazimine for 10 min, then treated with 1:10 diluted VSV-G or Spike for 10 min. Images were taken every 15 min. P, platelets; V, VSV-G; S, Spike; N, Niclosamide; C, Clofazimine. **B-C.** Graphs showing the percentage of clot retraction over a 90 min observation, upon PRP treatment with either PBS or VSV-G (B and C respectively) in the presence or absence of Niclosamide. Results are from n=4 independent experiments. Data are mean ± SEM. **D-F.** Same as panels A-C using Clofazimine.

